# Prefrontal-medullary circuitry is necessary for sex-specific responses to metabolic stress in rats

**DOI:** 10.64898/2026.02.10.705203

**Authors:** Carley Dearing, Ema Lukinic, Carlie McCartney, Brent Myers

## Abstract

Chronic stress increases risk for metabolic disorders, including diabetes mellitus.

Additionally, projections from the infralimbic cortex (IL) to the rostral ventrolateral medulla (RVLM) regulate endocrine stress responses. However, the neurobiological basis for chronic stress effects on glucose homeostasis has not been identified. The current study tests the hypothesis that the IL-RVLM circuit is necessary to prevent glucose intolerance. Accordingly, male and female rats with Cre-dependent expression of tetanus toxin light chain (TeLC) to inhibit neurotransmitter release from RVLM-projecting IL neurons were subject to chronic variable stress (CVS) or remained as No CVS controls. Animals were then acutely challenged with a fasted intraperitoneal glucose tolerance test (GTT). Endocrine metabolic function was evaluated during GTT via time courses of glucose, insulin, glucagon, and corticosterone. In No CVS females expressing TeLC, inhibition of IL-RVLM circuit signaling impaired glucose tolerance characterized by elevated glucose and decreased insulin sensitivity. Following chronic stress, females had impaired glucoregulation characterized by decreased glucose clearance and elevated corticosterone. When combined with TeLC, chronically-stressed females showed shifts in the ratio of insulin to glucagon compared to CVS GFP females, suggesting circuit function impacts the pancreatic mechanisms mediating glucose homeostasis during chronic stress. In No CVS males, TeLC increased glucagon only. However, CVS TeLC males had impaired glucose tolerance, reduced insulin sensitivity, and decreased corticosterone. These data indicate that the IL-RVLM circuit mediates glucoregulation in a manner dependent on both sex and stress history. Collectively, the IL-RVLM circuit is necessary for the sex-specific maintenance of glucose homeostasis following chronic stress.

## 1. Introduction

Metabolic disorders, such as diabetes mellitus, increasingly impact mortality and years lived with disability (1). However, a larger population experiences symptoms of metabolic dysfunction, referred to as metabolic syndrome, at a subclinical level (2). Symptoms of metabolic syndrome increase subsequent disease risk and are characterized by abdominal obesity, insulin resistance, hypertension, and hyperlipidemia (3). While the etiology of metabolic disorders and metabolic syndrome are multifactorial (2,4,5), environmental factors, such as stress, strongly contribute to the pathogenesis of these conditions (6–8). Broadly defined, stress is a real or perceived threat to homeostasis or well-being and results from a variety of life events (9,10).

Cumulative life stressors reduce organismal adaptive capacity and increase susceptibility to pathology (11). Importantly, stress-related conditions have sex-specific profiles that are important for both prognosis and treatment outcomes (12,13).

Exposure to chronic stress associates with deleterious changes across multiple physiological systems that impact metabolic function (8,14). Chronic stress can particularly impact cognitive and emotional processes and contribute to abdominal obesity and metabolic syndrome (6,15,16). Moreover, studies in rats found sexually divergent effects of chronic stress on glucose tolerance, as well as sex differences in the facilitation of glucocorticoid responses after chronic stress (17). Prolonged stress also increases the risk of hypertension and myocardial infarction (18,19), inflammation and chronic pain (20,21), as well as circadian disruption (22). Collectively, these changes promote homeostatic dysregulation and impact metabolic function and adaptive capacity.

While stress appraisal occurs in multiple brain regions, the human ventromedial prefrontal cortex (vmPFC) and rodent homolog, the infralimbic cortex (IL), are important for limiting male hypothalamic-pituitary-adrenal (HPA) axis responses (23) and influence coping behaviors after chronic stress (24). However, the IL has sex-specific impacts on socio-motivational behaviors and stress reactivity (25). Importantly, the IL does not directly innervate stress-regulatory neuroendocrine cells, indicating intermediary circuitry underlies cortical influences on stress responding (26,27). Activation of projections to the rostral ventrolateral medulla (RVLM), an important regulator of sympathetic activity and glucoregulation (28–33), was recently found to sex-specifically reduce stress responsivity (34).

Therefore, to test the necessity of the IL-RVLM circuit for glucoregulation following chronic variable stress (CVS) in males and females, we used an intersectional genetic approach to selectively reduce circuit signaling followed by glycemic challenge with an acute metabolic stress test. Specifically, we hypothesized that disruption of the IL-RVLM circuit would exacerbate the glucodysregulation associated with chronic stress where females show greater susceptibility to metabolic dysfunction (17).

## 2. Methods

### 2.1 Subjects

Age-matched young adult (PND ∼42) female and male Sprague-Dawley rats were obtained from Charles River (Wilmington, MA). Animals were kept on a 12-hour light cycle, lights on:off (07:00-19:00), with food and water *ad libitum*. Following stereotaxic surgery, rats were housed individually in a temperature-and humidity-controlled room. All experiments were approved by the Colorado State University Institutional Animal Care and Use Committee (Protocol:1392) and complied with the National Institutes of Health Guidelines for the Care and Use of Laboratory Animals. All rats had daily welfare checks by veterinary and/or animal resource staff.

### 2.2 Design

Staggered cohorts of young adult female and male rats (2 treatment-balanced cohorts/sex) were assigned to a 2X2 design with 4 groups per sex of either circuit-intact (green fluorescent protein [GFP]) or circuit-inhibited (tetanus toxin light chain [TeLC]) and No CVS or CVS. Animals were randomly assigned to both viral group (GFP or TeLC) and stress group (No CVS or CVS) within sex (n = 8-11/group/sex). CVS and No CVS groups within sex were weight-matched prior to CVS to prevent differences in starting body weight. Female and male cohorts were run consecutively, limiting between-sex analyses. CVS began 5 weeks after viral injection (D0-D14) (**Fig. 1A**). Fasted glucose tolerance testing (GTT) took place on day 15 followed by tissue collection on day 18.

**Figure 1.**
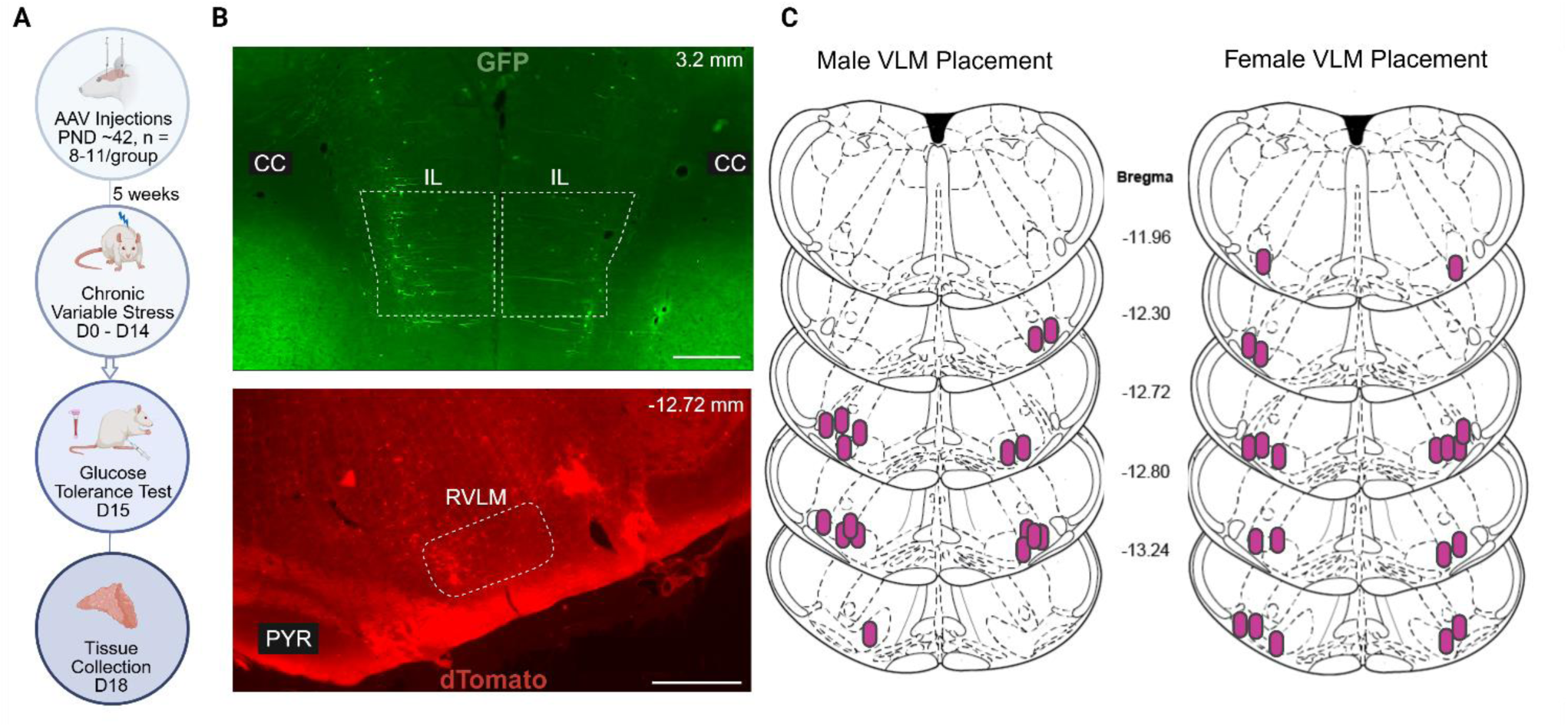
Experimental design and injection placement. Two distinct cohorts of male (n = 8-10/group) and female (n = 9-11/group) rats underwent viral injection followed by chronic stress exposure. Rats were then acutely challenged via GTT with blood and tissue collected (A). Representative micrographs (B) of viral injection placement in both IL (dashed box) [AAV-CMV-DIO-eGFP-TeLC] and RVLM (dashed box) [AAVrg-hSyn-Cre-dTomato]. Only males and females (C) with verified on-site injections and intersectional recombination were included in analyses. Created with BioRender. CC: corpus callosum, PYR: pyramidal tract

### 2.3 Microinjections

An intersectional genetic approach was used to reduce neurotransmitter release from RVLM-projecting IL neurons. Animals were first anesthetized with isoflurane (1-5%) followed by subcutaneous administration of analgesic buprenorphine-SR (0.6 mg/kg). Unilateral RVLM injections (female: 12.25 mm posterior to bregma, 1.9 mm lateral to midline, 9.95 mm ventral from skull; male: 12.25 mm posterior to bregma, 1.9 mm lateral to midline, 10.2 mm ventral from skull) were administered with a retrograde-transported adeno-associated virus (AAV; female: 0.3 μl, male: 0.5 μl; Addgene, Watertown, MA) carrying a Cre recombinase construct (hSyn-Cre-dTomato). Injection lateralization was randomized and balanced within and across groups. Although unilateral targeting may reduce the extent of Cre expression, the approach limited respiratory disruptions associated with bilateral RVLM injections. Subsequently, bilateral IL injections (female: 2.3 mm anterior to bregma, 0.5 mm lateral to midline, and 4 mm ventral from dura; male: 2.7 mm anterior to bregma, 0.6 mm lateral to midline, and 4.2 mm ventral from dura) were administered (female: 0.6 μl, male: 0.8 μl) of either a Cre-dependent TeLC (CMV-DIO-GFP-TeLC; Stanford Gene Vector and Virus Core, Stanford, CA) or Cre-dependent GFP (CMV-DIO-GFP; Addgene, Watertown, MA) (**Fig. 1B**). TeLC cleaves synaptobrevin and inhibits neurotransmitter exocytosis, thereby reducing signaling in the IL-RVLM circuit (35–37). Surgical recovery and construct expression continued for 5 weeks prior to initiation of stressors.

### 2.4 Chronic stress paradigm

Five weeks following viral injections, rats began the 14-day CVS paradigm which consisted of twice daily (AM and PM) randomized stressors and interspersed overnight stressors (38–40). Stressors included restraint (plexiglass tube, 1 h), shaker (100 rpm, 1 h), cold room (4°C, 1 h), brightly lit open field (1 m^2^, 1 h), forced swim (23-27°C, 10 min), cage tilt (45°, 1 h), predator odor (fox or coyote urine, 1 h), PM light (overnight, 12 h), and damp bedding (overnight, 12 h). All rats were fed *ad libitum* throughout CVS with food intake measured twice weekly to determine average daily food intake. Food was removed the morning of day 15 for a 4-h fast prior to GTT.

### 2.5 Glucose tolerance

On the morning of day 15, glucose metabolism and endocrine stress reactivity were assessed using a 4-h fasted intraperitoneal GTT (17,41). GTT was conducted more than 12 h following the final stressor and took place between hours 3 - 6 of the light phase. A baseline blood sample was taken via tail clip prior to glucose injection (1.5 g/kg, 25% glucose, i.p.). Blood was then collected at 15, 30, 45, and 90 min post-injection and placed in tubes with 10 µl of 100 mmol/L ethylenediamine tetraacetate. Blood glucose was quantified in duplicate with Bayer Contour Next Glucometers and test strips (Ascensia, Parsippany, NJ). Blood was then centrifuged at 3000 x g for 15 min at 4° C and stored at-20° C until analysis. Plasma corticosterone (C. V. = 6.6 – 8.0 %; ENZO Life Sciences, Farmingdale, NY), plasma glucagon (C. V. < 10 %; Crystal Chem, Elk Grove Village, IL), and plasma insulin (C. V. ≤ 10 %; Crystal Chem, Elk Grove Village, IL) were determined by ELISA according to manufacturers’ instructions. Vaginal swabs were taken for estrous analysis in females following blood sampling.

### 2.6 Tissue collection and processing

Two days after the GTT, animals were deeply anesthetized with sodium pentobarbital (100 mg/kg i.p.) and perfused transcardially with 0.9% saline followed by 4.0% paraformaldehyde in 0.1 M phosphate buffer solution. Adrenal glands were collected and weighed and vaginal swabs were taken for estrous analysis in females. Brain tissue was collected for technical validation (**Fig. 1C**). Briefly, tissue was post-fixed in 4.0% paraformaldehyde for 24 hours at room temperature before storage in 30% sucrose at 4° C. Coronal sections were made on a freezing microtome at 30 μm thickness and then stored in cryoprotectant solution at −20° C. To verify injection and placement, coronal brain sections were imaged with a Zeiss Axio Imager Z2 microscope (Carl Zeiss Microscopy, Jena, Germany) using a 10x objective. Placement of retrograde AAV in RVLM was confirmed, as well as Cre-dependent reporter expression in IL. Cases with injection placement outside the RVLM were omitted from analyses.

### 2.7 Statistical analyses

Data are presented as mean ± standard error of the mean (SEM). Animals were excluded from all analyses if injection placement was greater than 0.5 mm from targeted region. All data were analyzed using Prism 10 (GraphPad, San Diego, CA) with significance defined as p < 0.05. All analyses were completed within sex. Body weight during CVS was analyzed by 3-way repeated measures ANOVA with stress (No CVS or CVS), virus (GFP or TeLC), and time (repeated) as factors. Food intake and adrenal weight were compared via 2-way ANOVA with stress and viral factors. Due to the neurocircuit-focused hypothesis, hormonal responses to GTT were analyzed via 2-way ANOVA within-stress with virus and time (repeated) as factors. Homeostatic model assessment of insulin resistance (HOMA-IR) was calculated on fasting parameters (42–44) and compared by 2-way ANOVA with stress and viral factors. In the case of ANOVA significant main or interaction effects, Tukey or Fisher (for repeated measures) post-hoc analysis was used for group comparisons. Correlative measures between glucose and either insulin, glucagon, or corticosterone were determined by Pearson’s r two-tailed analyses. All ANOVA results are included in supplemental table 1 (**Table S.1**).

## 3. Results

### 3.1 Body weight, food intake, and adrenal index

To examine somatic markers of chronic stress, body weight, food intake, and adrenal gland size were assessed. In females (n = 8-10/group), body weight analysis over the course of CVS showed main effects of stress [F(1, 33) = 6.641, p = 0.0146] and time [F(4, 132) = 6.825, p < 0.0001] and an interaction of stress and time [F(4, 132) = 23.80, p < 0.0001] where chronically-stressed females had reduced body weight compared to No CVS rats, regardless of viral treatment (**Fig. 2A**). Additionally, mean daily food intake had a main effect of stress [F(1, 33) = 13.58, p = 0.0008] where CVS TeLC females had significantly less food intake than No CVS TeLC females (p = 0.0402) (**Fig. 2B**). However, there were no significant differences in body weight-corrected adrenal weight in females (**Fig. 2C**).

**Figure 2.**
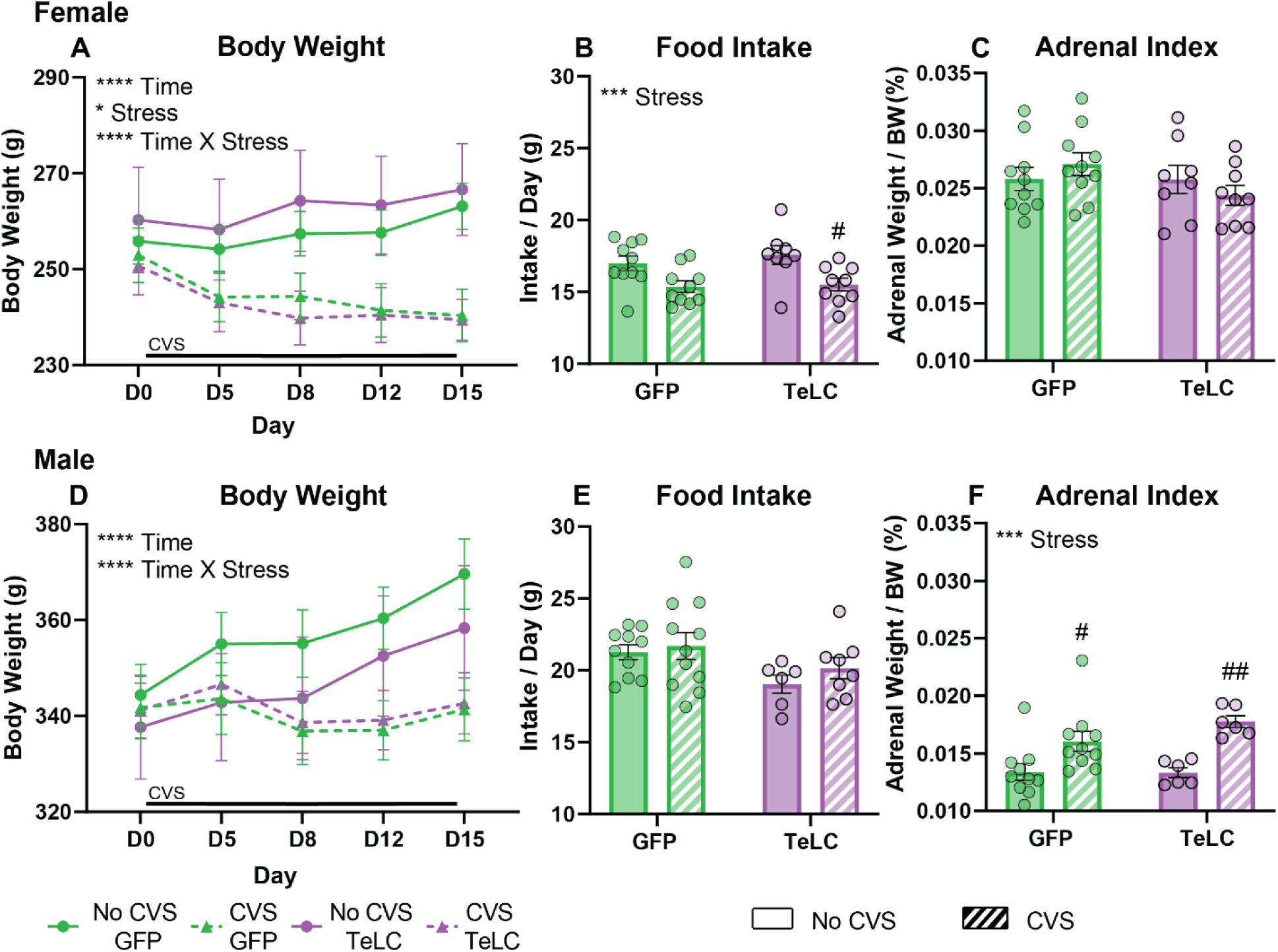
Somatic measures following CVS. Female (n = 9-11/group) body weight (A), food intake (B), and adrenal weight corrected for body weight (C) were measured over the course of CVS. Male (n = 8-10/group) body weight (D), food intake (E), and adrenal index (F) were analyzed. Data are expressed as mean ± SEM. * p<0.05, *** p<0.001, **** p<0.0001 ANOVA effects; ^#^ p < 0.05, ^##^ p < 0.01 CVS vs. No CVS.

In male rats (n = 9-11/group), body weight over the course of CVS had a main effect of time [F(4, 124) = 44.01, p < 0.0001] and an interaction of stress and time [F(4, 124) = 49.58, p < 0.0001] (**Fig. 2D**) where CVS-exposed animals had lower body weight than No CVS, independent of circuit manipulation. There were no significant differences in male food intake over CVS (**Fig. 2E**). However, there was a main effect of stress on male adrenal weight [F(1, 28) = 20.32, p = 0.0001] where both CVS GFP (p = 0.0469) and CVS TeLC (p = 0.0069) groups had significantly increased relative adrenal weight than No CVS counterparts (**Fig. 2F**).

### 3.2 Hormonal responses to GTT

Endocrine responses to acute metabolic stress were determined via glycemic challenge. In females (n = 8-10/group), No CVS TeLC animals had decreased glucose clearance [F(1, 79) = 10.36, p = 0.0019] at 15 (p = 0.0042) and 30 (p = 0.0079) min (**Fig. 3A**), compared to No CVS GFP controls. Interestingly, TeLC did not significantly affect female plasma insulin or glucagon responses to hyperglycemia (**Fig. 3B, C**). However, TeLC increased corticosterone secretion [F(1, 79) = 14.41, p = 0.0003] 30 (p = 0.044) and 45 (p = 0.0002) min post glucose injection (**Fig. 3D**).

**Figure 3.**
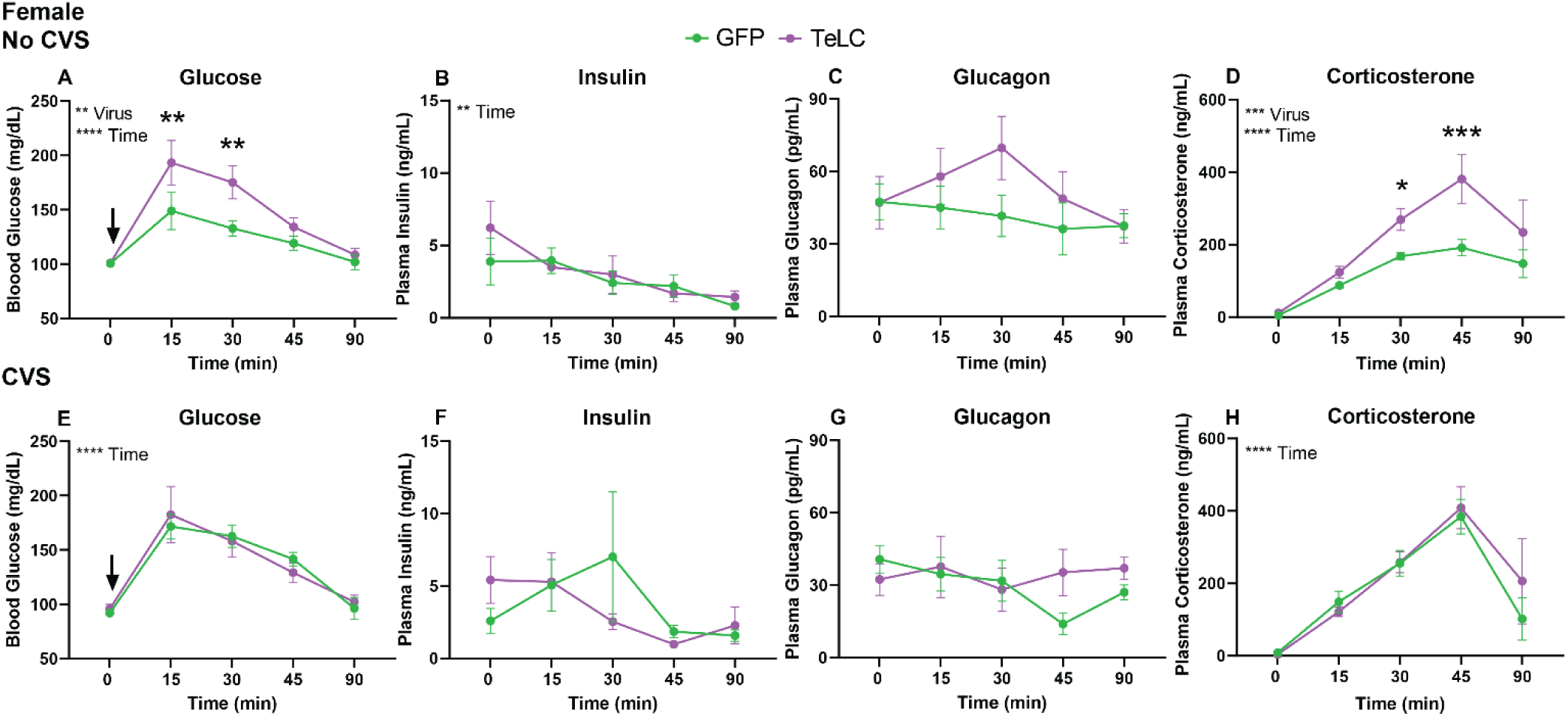
Female GTT hormonal responses. Following CVS, females were challenged with GTT (n = 8-10/group/sex). Animals were compared within stress history: No CVS GFP (n = 10), No CVS TeLC (n = 8), CVS GFP (n= 10), CVS TeLC (n = 9). Blood was collected at 0, 15, 30, 45, and 90 min. Blood glucose (A, E), plasma insulin (B, F), plasma glucagon (C, G), and plasma corticosterone (D, H) were quantified over the 90-minute time course. Data are expressed as mean ± SEM. ↓ represents i.p. glucose bolus. * p<0.05, ** p<0.01, *** p<0.001,**** p<0.0001 ANOVA effects or TeLC vs. GFP.

As our prior reports found that CVS in females both impairs glucose tolerance and facilitates corticosterone responses to glycemic stress (17), the current analysis focused on within-CVS effects of TeLC. Here, CVS TeLC females had no significant differences from CVS GFP females in terms of glucose, insulin, glucagon, or corticosterone responses (**Fig. 3E, F, G, H**); although, both GFP and TeLC females exposed to CVS had glucodysregulation evidenced by elevated glucose and corticosterone. Thus, the effects of female IL-RVLM inhibition do not interact with those of CVS to further exacerbate metabolic hormone time courses after chronic stress.

In males (n = 9-11/group), No CVS circuit inhibition did not overtly change glucose tolerance or insulin responses to GTT (**Fig. 4A, B**). However, these animals did have differences in underlying metabolic regulation that are reflected in alpha cell responsivity as No CVS TeLC males showed significantly increased glucagon compared to GFP controls [F(1, 63) = 4.015, p = 0.0494] (**Fig. 4C**). There were no significant differences in plasma corticosterone across the time course (**Fig. 4D**).

**Figure 4.**
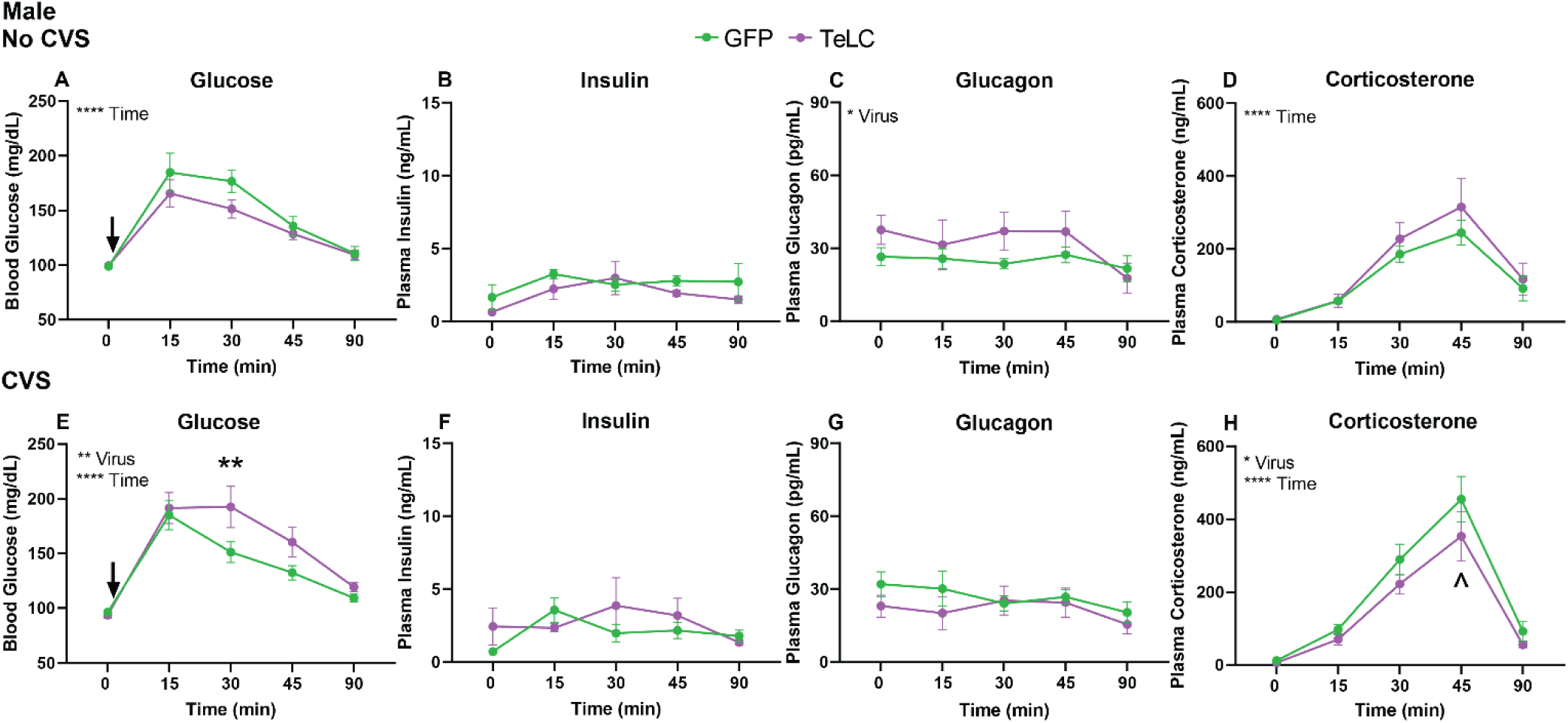
Male GTT hormonal responses. Following CVS, males were challenged with GTT (n = 6-11/group). Animals were compared within stress history: No CVS GFP (n = 10), No CVS TeLC (n = 6), CVS GFP (n= 11), CVS TeLC (n = 7). Blood was taken at 0, 15, 30, 45, and 90 min. Blood glucose (A, E), plasma insulin (B, F), plasma glucagon (C, G), and plasma corticosterone (D, H) were quantified over the 90-minute time course. Data are expressed as mean ± SEM. ↓ represents i.p. glucose bolus. * p<0.05, ** p<0.01, **** p<0.0001 ANOVA effects or TeLC vs. GFP; ^ p = 0.058 TeLC vs. GFP.

Following chronic stress, TeLC treatment in males led to deviations from the metabolic profile previously reported following CVS where chronic stress improves male glucose tolerance (17,45). Here, CVS TeLC males had decreased glucose clearance [F(1, 81) = 7.018, p = 0.0097] (**Fig. 4E**) 30 min after the glucose bolus (p = 0.0044). There were no significant differences in insulin or glucagon across the time course (**Fig. 4F, G**). However, TeLC decreased corticosterone [F(1, 80) = 4.078, p = 0.0468] (**Fig. 4H**), most evident at 45 min (p = 0.058). Taken together, the necessity of the IL-RVLM circuit for regulating metabolic hormonal responses to hyperglycemia is sex-and stress history-specific, further indicating that both biological and environmental factors interact to regulate glucose homeostasis.

### 3.3 Metabolic integration and insulin sensitivity

To examine the integration of endocrine and autonomic contributions to metabolic regulation, relative pancreatic islet cell responsivity was examined by calculating the ratio of plasma insulin to plasma glucagon (I:G). This measure queries the relative responsivity of alpha cells (sympathetic) and beta cells (parasympathetic) in the pancreas resulting from both autonomic and HPA axis signals, which is vital to glucoregulation and homeostatic adaptation (46–48). In females, there were no significant differences in I:G between the No CVS GFP and TeLC groups (**Fig. 5A)**. However, following CVS, females had an interaction of virus and time on I:G [F(4, 77) = 2.57, p = 0.0444] where the CVS GFP females had greater relative insulin responsivity and CVS TeLC females shifted toward greater glucagon reactivity (**Fig. 5B**) 45 min following glucose administration (p = 0.0102). Analysis of fasting baseline insulin and glucose was carried out according to the HOMA-IR formula where elevated values indicate decreased insulin sensitivity and pathological insulin resistance is typically defined by values above 2 (42–44). This analysis (**Fig. 5C**) indicated that TeLC treatment impaired insulin sensitivity in females regardless of stress condition [F(1, 31) = 9.867, p = 0.0037], suggesting that these animals may carry susceptibility to metabolic dysregulation.

**Figure 5.**
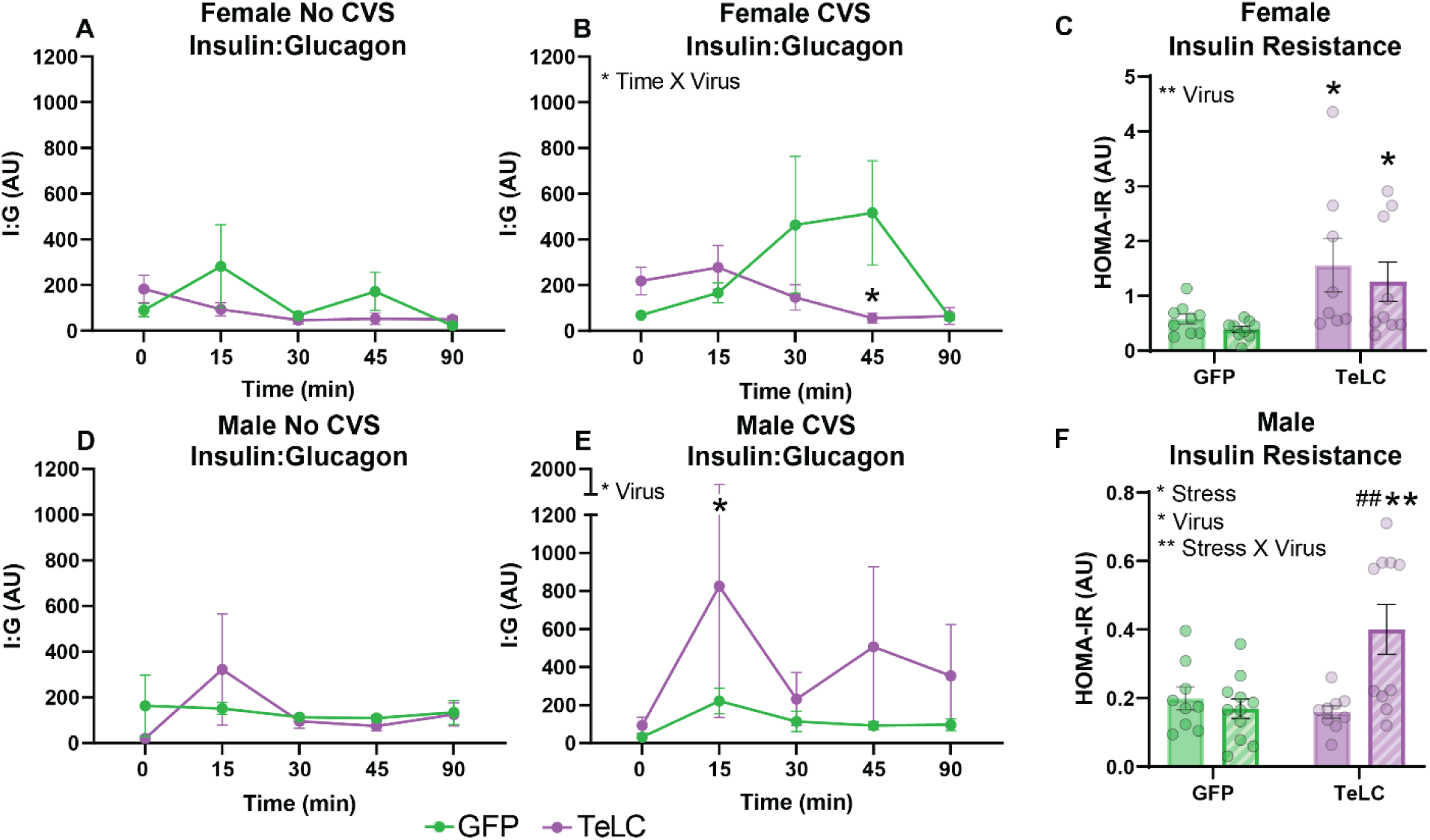
Insulin:glucagon over GTT and fasting insulin resistance. The ratio of I:G was quantified at each time point for both females (A, B) and males (D, E) by stress history (n = 6-11/group/sex). Homeostatic model assessment of insulin resistance (HOMA-IR) was evaluated in fasting animals (C, F). Data are expressed as mean ± SEM. * p<0.05, ** p<0.01, ANOVA effects or TeLC vs. GFP; ^##^ p < 0.01 vs. No CVS.

In males without prior stress exposure, there were no effects of circuit manipulation on pancreatic reactivity (**Fig. 5D**). Although, TeLC males exposed to chronic stress had increased I:G compared to GFP animals [F(1, 76) = 5.048, p = 0.0276] (**Fig. 5E**) 15 min post glucose (p = 0.0445), indicating greater beta cell responsivity to hyperglycemia. Similarly, TeLC impaired fasting baseline insulin sensitivity only in males exposed to CVS (**Fig. 5F**). Here, there were main effects of stress [F(1, 35) = 5.687, p = 0.0226], virus [F(1, 35) = 4.653, p = 0.0379], and a virus by stress interaction [F(1, 35) = 9.362, p = 0.0042] where CVS TeLC males had impaired insulin sensitivity relative to all other groups (p < 0.01).

### 3.4 Endocrine associations with glucose tolerance

Regressive correlations were used to examine whether glucose levels associated with regulatory hormones over the course of the GTT among the individual subjects in each group. These analyses highlight relationships between glucoregulatory hormones that may account for individual variability and/or group-specific outcomes. In No CVS GFP females, 90-min plasma insulin negatively correlated with blood glucose at multiple times post-injection (15 min: r = - 0.92, p = 0.009; 45 min: r = - 0.896, p = 0.016; 90 min: r = - 0.85, p = 0.032) (**Fig. 6A**).

**Figure 6.**
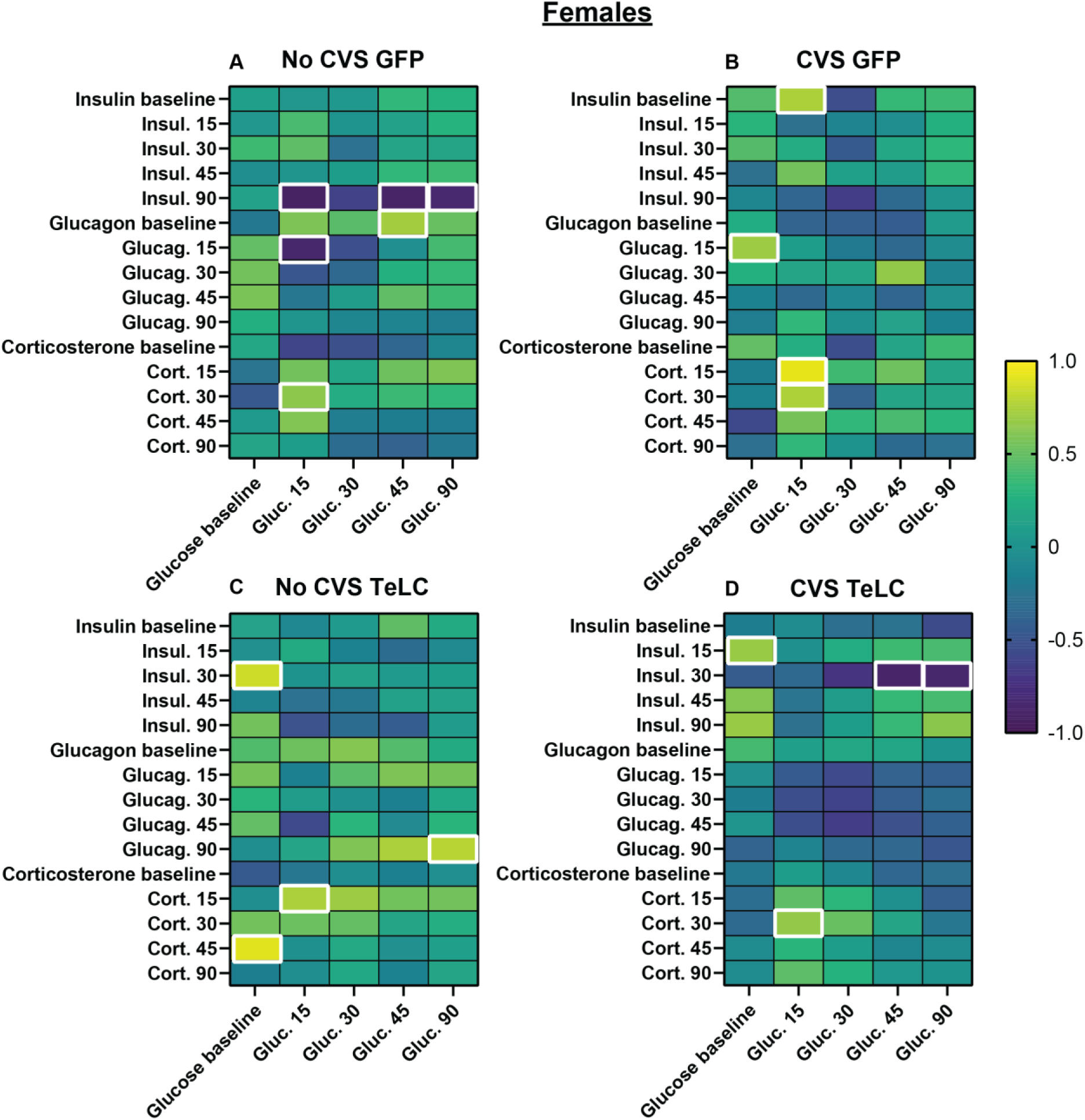
Regressive analysis: females. Pearson correlations were run within each group to examine the relationship between glucose and insulin, glucagon, and corticosterone. Significant associations (p < 0.05) are bolded.

Alternatively, 15-min blood glucose positively correlated with corticosterone (30 min: r = 0.644, p = 0.044), and 45-min glucose positively correlated with baseline glucagon (r = 0.72, p = 0.019). Taken together, these relationships in female controls indicate systems integration where gluconeogenic hormones associate with elevated glucose and insulin with reduced glucose.

Following CVS, there were shifts in these associations. Specifically, chronically-stressed GFP-treated females had no significant negative correlations between insulin and glucose. Instead, baseline glucose positively related to glucagon (15 min: r = 0.699, p = 0.024) (**Fig. 6B**) and the initial rise in glucose associated with elevated corticosterone (15 min: r = 0.927, p < 0.0001; 30 min: r = 0.756, p = 0.018). Surprisingly, baseline insulin positively correlated with blood glucose (15 min: r = 0.75, p = 0.032), indicating potential disruption of fundamental glucoregulatory processes in females after chronic stress.

The knockdown of IL-RVLM signaling in No CVS females led to positive correlations of baseline glucose with subsequent insulin (30 min: r = 0.853, p = 0.007) and corticosterone (45 min: r = 0.908, p = 0.002) responses to hyperglycemia (**Fig. 6C**). Additional time-matched positive associations were present for glucose with corticosterone (15 min: r = 0.745, p = 0.034) and glucagon (90 min: r = 0.791, p = 0.034). Altogether, No CVS TeLC females had decreases in both glucose tolerance and insulin sensitivity which may relate to the lack of significant negative relationships between glucose and insulin.

Examining endocrine relationships in CVS females with IL-RVLM circuit knockdown revealed a unique pattern of hormonal associations. As in other female groups, there was a positive relationship between glucose and corticosterone (30 min: r = 0.666, p = 0.049) (**Fig. 6D**). However, unlike either CVS or TeLC alone, circuit inhibition after chronic stress led to negative correlations between 30-min insulin and subsequent glucose (45 min: r =-0.886, p = 0.008; 90 min: r =-0.854, p = 0.014), potentially pointing to individual variability of insulin sensitivity within the group.

Overall, there were fewer significant endocrine correlations across the male groups. In No CVS GFP male rats, baseline glucose positively correlated with 45-min glucagon (r = 0.79, p =0.021) and insulin (r = 0.71, p = 0.031). Additionally, baseline glucagon and 45-min glucagon both correlated with subsequent glucose at (30 min: r = 0.61, p = 0.031; 90 min: r = 0.74, p = 0.036) (**Fig. 7A**), suggesting that glucose most closely associates with glucagon substrate in control males. Interestingly, chronically-stressed GFP males had no significant correlations of glucose with either insulin, glucagon, or corticosterone (**Fig. 7B**). However, No CVS TeLC males had a single positive correlation between 15-min insulin and glucose at 30 min (r = 0.83, p = 0.04) (**Fig. 7C**). Unique relative to the other male groups, TeLC CVS males had glucose intolerance and insulin insensitivity, coupled with reduced corticosterone secretion. Examination of hormonal interactions indicated that baseline glucose negatively correlated with baseline glucagon (r =-0.08, p = 0.032) and 45-min glucagon (r =-0.81, p = 0.026) (**Fig. 7D**). These negative associations between glucagon and glucose along with the absence of significant relationships between glucose and either insulin or corticosterone suggest that CVS TeLC males have alterations in the primary bases of metabolic function.

**Figure 7.**
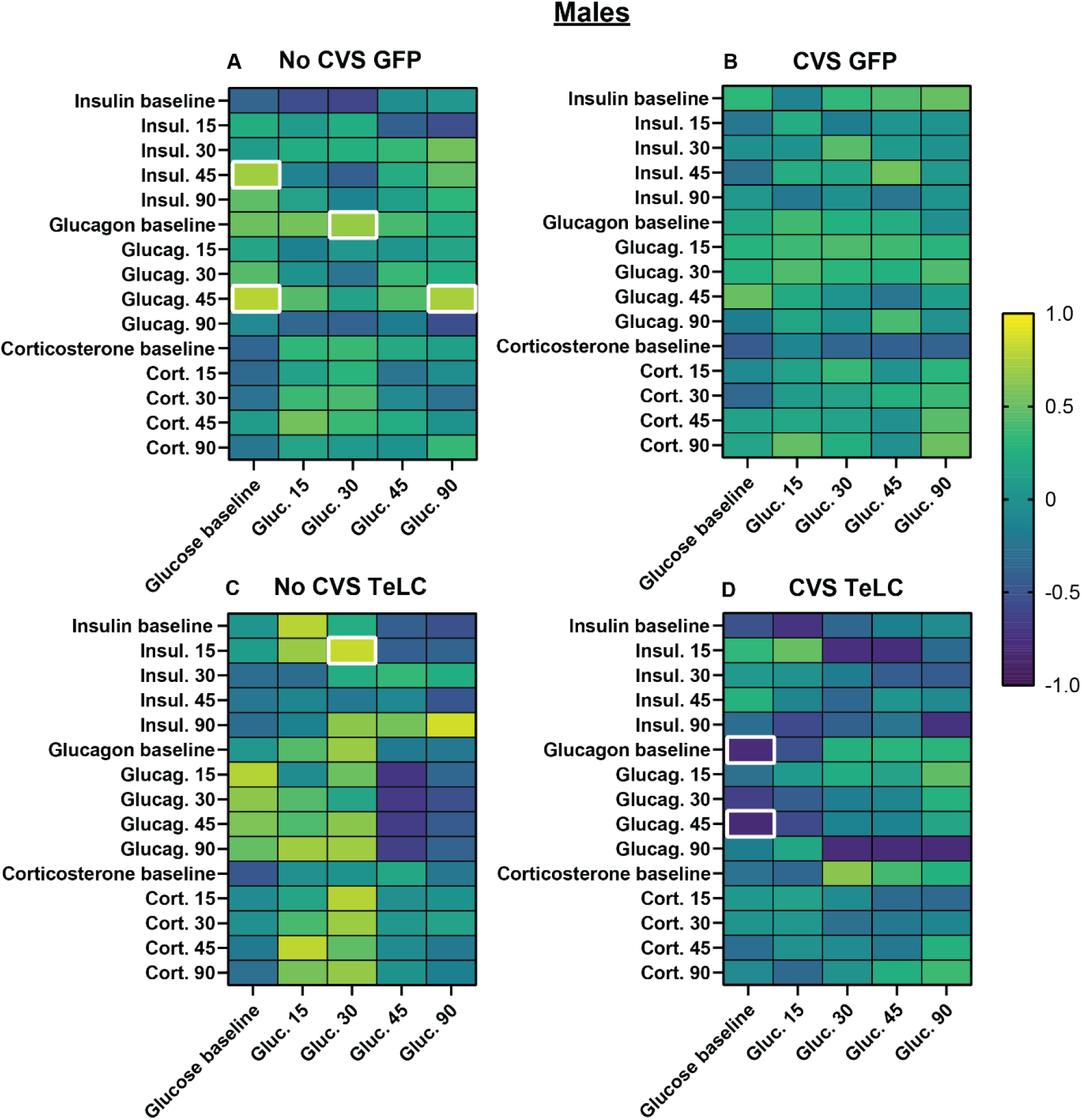
Regressive analysis: males. Pearson correlations were run within each group to examine the relationship between glucose and insulin, glucagon, and corticosterone. Significant associations (p < 0.05) are bolded.

### 3.5 Estrous cycle

Vaginal cytology was used to determine estrous phase following GTT and at tissue collection (**Table 1**). All rats were free cycling and were not staged to allow for representation of the female reproductive profile, generally. Sample size per phase was not sufficient to permit cycle-specific analysis but prior work with large sample sizes found no significant relationships between estrous phase and glucose tolerance (17). Here, the proportional representation of each phase is reported with cycle variability present across assessments.

**Table 1:** Estrous stage at assessment (%).

|  | GTT |  |  |  | Tissue Collection |  |  |  |
| --- | --- | --- | --- | --- | --- | --- | --- | --- |
|  | Proestrus | Estrus | Metestrus | Diestrus | Proestrus | Estrus | Metestrus | Diestrus |
| **No**<br>**CVS**<br>**GFP** | 20 | 20 | 10 | 50 | 0 | 30 | 40 | 30 |
| **CVS**<br>**GFP** | 40 | 20 | 40 | 0 | 10 | 50 | 20 | 20 |
| **No**<br>**CVS**<br>**TeLC** | 63.6 | 9.1 | 27.3 | 0 | 9.1 | 54.5 | 9.1 | 27.3 |
| **CVS**<br>**TeLC** | 54.5 | 18.2 | 18.2 | 9.1 | 18.2 | 27.3 | 36.4 | 18.2 |

## 4. Discussion

### 4.1 IL-RVLM glucose regulation: females

Glucoregulation is a complex and dynamic process that is impacted by sex and prior stress history (17,30,32,33,45,49,50). The current results indicate that IL-RVLM circuitry is necessary for preventing glucose intolerance and insulin insensitivity in previously unstressed females.

Specifically, circuit inhibition reduced fasting insulin sensitivity, as well as impairing glucose clearance and elevating corticosterone responses to hyperglycemia. Altogether, suggesting that the female IL-RVLM circuit acts to tonically inhibit sympathetic and HPA axis reactivity.

Knocking down signaling in the circuit led to physiological drive characterized by associations with gluconeogenic hormones, including correlations of glucose with both glucagon and corticosterone. Ultimately, impairments to signaling of IL synapses in the female RVLM promote homeostatic dysfunction which may generate a state of metabolic susceptibility.

Although chronic stress impairs female glucose clearance and increases corticosterone release (17), females undergoing chronic stress with knockdown of IL-RVLM signaling did not have further exacerbations of glucose intolerance or corticosterone hypersecretion. However, CVS TeLC females had decreased fasted insulin sensitivity and a shift toward alpha cell reactivity during hyperglycemia relative to chronically-stressed GFP females, indicting an increase in gluconeogenic processes. Therefore, chronic stress in females promotes dysregulation of glucose homeostasis with circuit knockdown further disrupting insulin sensitivity.

### 4.2 IL-RVLM glucose regulation: males

In male rats, the impact of inhibiting the IL-RVLM circuit was limited. Unstressed animals did not have glucose intolerance or corticosterone hypersecretion. However, these animals did exhibit increased glucagon throughout the GTT, indicating a potential facilitation of gluconeogenesis even in the context of hyperglycemia. This increase in gluconeogenesis may reflect an increase in net sympathetic activation, similar to their female counterparts. However, the lack of corresponding changes in glucose tolerance could be indicative of increased male metabolic adaptive capacity. Although chronic stress in male rats increases corticosterone release, male glucose homeostasis improves after CVS (17,45). However, following exposure to chronic stress, male IL-RVLM inhibition decreased glucose clearance and insulin sensitivity which was underscored by a significant shift toward beta cell reactivity. Interestingly, these animals had decreased corticosterone secretion suggesting that reducing circuit signaling may disassociate autonomic and HPA axis regulation during chronic stress, an outcome supported by the absence of correlation between corticosterone and glucose in this group. The interplay between neuroendocrine processes and autonomic balance in CVS TeLC males suggests they may compensate for impaired circuit function and stress exposure differently than females.

### 4.3 Sex-dependent, stress-modulated neurocircuit control of homeostatic function

The regulation of hormone levels as well as the relationships between glucoregulatory hormones varied substantially based on sex, stress history, and circuit disruption. The findings in aggregate highlight the necessity of cortical-medullary circuit function for glucose homeostasis in unstressed females, while mediating aspects of chronic stress susceptibility in males. Prior studies of IL-VLM circuit organization found that both males and females have similar proportions of stress-responsive neurons projecting to the VLM (34). Both sexes also have widespread inputs to the VLM with IL glutamate neurons innervating the extent of the RVLM and caudal VLM (34). IL synapses contact epinephrine/norepinephrine, GABA, and glycine cells (34,51), indicating the capacity for context-specific regulation of the VLM network. While there may be sex-biased circuit signaling, it is unclear how circuit function interacts with chronic stress divergently in males and females. These effects may arise from a combination of genetic and developmental mechanisms and/or activational effects of gonadal hormones (50,52–54). Gonadal hormone receptors are expressed widely throughout central and peripheral tissues and contribute to glucose regulation (12,50,55). The current female results were not specific to a particular estrous stage and design did not permit analysis of potential cycle-dependent effects. However, estradiol and progesterone have divergent effects on glucose tolerance and insulin sensitivity. Estradiol enhances insulin sensitivity and reduces hepatic glucose production (56,57), while progesterone can increase insulin resistance (58). Androgenic regulation of glucose tolerance is also multi-dimensional, yet testosterone generally promotes male metabolic homeostasis and insulin sensitivity (56,59). Moreover, sex-specific chronic stress-responsiveness of transcriptional and epigenetic mechanisms in brain, adrenal, pancreas, liver, or other metabolic organs (60–64) could also contribute to divergent outcomes.

### 4.4 Conclusion

The current study examined the necessity of the IL-RVLM circuit for glucose homeostasis and metabolic stress responses following chronic stress exposure. The resulting data indicate that the IL-RVLM circuit mediates context-specific metabolic homeostasis as inhibition of the circuit produced sex-dependent glucodysregulation based on stress history. While these experiments were analyzed within sex, future work directly comparing male and female outcomes may elucidate sex differences in metabolic dysfunction after chronic stress.

Additionally, further examination of insulin signaling may reveal downstream mechanisms of metabolic dysfunction. Overall, this study provides novel insight into the glucoregulatory function of cortical-medullary circuitry and how adverse experience interacts in a sex-specific manner to tune metabolic susceptibility/resilience. Ultimately, altered function of the circuit may contribute to the health burden of stress and represent a novel target for intervention in metabolic disorders.

## 5. Acknowledgements

The authors would like to thank Sebastian Pace, Derek Schaeuble and Tyler Wallace for help with sample collection and Payton Shaffer for aid in tissue processing. The Cre-dependent GFP-expressing construct was a gift from Dr. Bryan Roth (Addgene viral prep no. 50465-AAV5) and the retrograde Cre was a gift from Dr. Rylan Larsen (Addgene viral prep no. 107738-AAVrg).

## Supplementary Material

**Table S1:** ANOVA table of main and interaction effects. All ANOVA results from all comparisons within the study.

|  | ANOVA Table | F(DFn,DFd) | P Value |
| --- | --- | --- | --- |
| **Figure 2** |  |  |  |
| Female body weight | Time | F (4, 132) = 6.825 | $P<0.0001$ |
| | Virus | F (1, 33) = 0.05442 | $P=0.8170$ |
| | Stress | F (1, 33) = 6.641 | $P=0.0146$ |
| | Time x Virus | F (4, 132) = 0.1295 | $P=0.9714$ |
| | Time x Stress | F (4, 132) = 23.80 | $P<0.0001$ |
| | Virus x Stress | F (1, 33) = 0.2910 | $P=0.5932$ |
| | Time x Virus x Stress | F (4, 132) = 0.8380 | $P=0.5034$ |
| Female food intake | Interaction | F (1, 33) = 0.1905 | $P=0.6653$ |
| | Virus | F (1, 33) = 0.5219 | $P=0.4751$ |
| | Stress | F (1, 33) = 13.58 | $P=0.0008$ |
| Female adrenal glands | Interaction | F (1, 33) = 1.712 | $P=0.1998$ |
| | Virus | F (1, 33) = 1.829 | $P=0.1854$ |
| | Stress | F (1, 33) = 0.002132 | $P=0.9634$ |
| Male body weight | Time | F (4, 124) = 44.01 | $P<0.0001$ |
| | Virus | F (1, 31) = 0.2922 | $P=0.5927$ |
| | Stress | F (1, 31) = 2.043 | $P=0.1630$ |
| | Time x Virus | F (4, 124) = 0.4410 | $P=0.7787$ |
| | Time x Stress | F (4, 124) = 49.58 | $P<0.0001$ |
| | Virus x Stress | F (1, 31) = 0.5418 | $P=0.4672$ |
| | Time x Virus x Stress | F (4, 124) = 1.680 | $P=0.1588$ |
| Male food intake | Interaction | F (1, 31) = 0.1795 | $P=0.6748$ |
| | Virus | F (1, 31) = 5.539 | $P=0.251$ |
| | Stress | F (1, 31) = 0.9533 | $P=0.3364$ |
| Male adrenal glands | Interaction | F (1, 28) = 1.273 | $P=0.2688$ |
| | Virus | F (1, 28) = 1.112 | $P=0.3007$ |
| | Stress | F (1, 28) = 20.32 | $P=0.0001$ |
| **Figure 3** |  |  |  |
| Female No CVS glucose | Interaction | F (4, 79) = 1.816 | $P=0.1340$ |
| | Time | F (4, 79) = 16.47 | $P<0.0001$ |
| | Virus | F (1, 79) = 10.36 | $P=0.0019$ |
| Female No CVS insulin | Interaction | F (4, 77) = 0.6094 | $P=0.6571$ |
| | Time | F (4, 77) = 4.104 | $P=0.0046$ |
| | Virus | F (1, 77) = 0.6074 | $P=0.4381$ |
| Female No CVS glucagon | Interaction | F (4, 75) = 0.7674 | $P=0.5498$ |
| | Time | F (4, 75) = 1.103 | $P=0.3616$ |
| | Virus | F (1, 75) = 3.015 | $P=0.0866$ |
| Female No CVS corticosterone | Interaction | F (4, 79) = 2.042 | $P=0.0965$ |
| | Time | F (4, 79) = 19.18 | $P<0.0001$ |
| | Virus | F (1, 79) = 14.41 | $P=0.0003$ |
| Female CVS glucose | Interaction | F (4, 85) = 0.3202 | $P=0.8637$ |
| | Time | F (4, 85) = 20.13 | $P<0.0001$ |
| | Virus | $F(1, 85) = 0.008482$ | $P=0.9268$ |
| Female CVS insulin | Interaction | $F(4, 82) = 1.005$ | $P=0.4098$ |
| | Time | $F(4, 82) = 1.546$ | $P=0.1965$ |
| | Virus | $F(1, 82) = 0.06800$ | $P=0.7949$ |
| Female CVS glucagon | Interaction | $F(4, 80) = 1.183$ | $P=0.3246$ |
| | Time | $F(4, 80) = 0.8370$ | $P=0.5057$ |
| | Virus | $F(1, 80) = 0.8857$ | $P=0.3495$ |
| Female CVS corticosterone | Interaction | $F(4, 83) = 0.5061$ | $P=0.7313$ |
| | Time | $F(4, 83) = 17.21$ | $P<0.0001$ |
| | Virus | $F(1, 83) = 0.3916$ | $P=0.5332$ |
| **Figure 4** |  |  |  |
| Male No CVS glucose | Interaction | $F(4, 70) = 0.5888$ | $P=0.6718$ |
| | Time | $F(4, 70) = 19.82$ | $P<0.0001$ |
| | Virus | $F(1, 70) = 2.461$ | $P=0.1212$ |
| Male No CVS insulin | Interaction | $F(4, 67) = 0.4124$ | $P=0.7991$ |
| | Time | $F(4, 67) = 1.569$ | $P=0.1928$ |
| | Virus | $F(1, 67) = 2.308$ | $P=0.1334$ |
| Male No CVS glucagon | Interaction | $F(4, 63) = 0.6312$ | $P=0.6420$ |
| | Time | $F(4, 63) = 1.385$ | $P=0.2492$ |
| | Virus | $F(1, 63) = 4.015$ | $P=0.0494$ |
| Male No CVS corticosterone | Interaction | $F(4, 68) = 0.4077$ | $P=0.8025$ |
| | Time | $F(4, 68) = 23.67$ | $P<0.0001$ |
| | Virus | $F(1, 68) = 1.901$ | $P=0.1725$ |
| Male CVS glucose | Interaction | $F(4, 81) = 1.623$ | $P=0.1763$ |
| | Time | $F(4, 81) = 31.22$ | $P<0.0001$ |
| | Virus | $F(1, 81) = 7.018$ | $P=0.0097$ |
| Male CVS insulin | Interaction | $F(4, 78) = 1.356$ | $P=0.2570$ |
| | Time | $F(4, 78) = 1.431$ | $P=0.2315$ |
| | Virus | $F(1, 78) = 1.270$ | $P=0.2632$ |
| Male CVS glucagon | Interaction | $F(4, 78) = 0.3960$ | $P=0.8109$ |
| | Time | $F(4, 78) = 0.8988$ | $P=0.4689$ |
| | Virus | $F(1, 78) = 2.237$ | $P=0.1388$ |
| Male CVS corticosterone | Interaction | $F(4, 80) = 0.4972$ | $P=0.7378$ |
| | Time | $F(4, 80) = 37.56$ | $P<0.0001$ |
| | Virus | $F(1, 80) = 4.078$ | $P=0.0468$ |
| **Figure 5** |  |  |  |
| Female No CVS I:G | Interaction | $F(4, 74) = 1.197$ | $P=0.3192$ |
| | Time | $F(4, 74) = 1.351$ | $P=0.2594$ |
| | Virus | $F(1, 74) = 0.7814$ | $P=0.3796$ |
| Female CVS I:G | Interaction | $F(4, 77) = 2.570$ | $P=0.0444$ |
| | Time | $F(4, 77) = 1.415$ | $P=0.2370$ |
| | Virus | $F(1, 77) = 1.862$ | $P=0.1764$ |
| Female HOMA-IR | Interaction | $F(1, 31) = 0.0364$ | $P=0.8499$ |
| | Stress | $F(1, 31) = 0.6928$ | $P=0.4116$ |
| | Virus | $F(1, 31) = 9.867$ | $P=0.0037$ |
| Male No CVS I:G | Interaction | $F(4, 63) = 0.8112$ | $P=0.5227$ |
| | Time | $F(4, 63) = 0.9507$ | $P=0.4409$ |
| | Virus | $F(1, 63) = 0.01388$ | $P=0.9066$ |
| Male CVS I:G | Interaction | $F(4, 76) = 0.5922$ | $P=0.6693$ |
| | Time | $F(4, 76) = 1.414$ | $P=0.2373$ |
| | Virus | $F(1, 76) = 5.048$ | $P=0.0276$ |
| Male HOMA-IR | Interaction | $F(1, 35) = 9.362$ | $P=0.0042$ |
| | Stress | $F(1, 35) = 5.687$ | $P=0.0226$ |
| | Virus | $F(1, 35) = 4.653$ | $P=0.0379$ |

## References

1. Chew NWS, Ng CH, Tan DJH, Kong G, Lin C, Chin YH, Lim WH, Huang DQ, Quek J, Fu CE, Xiao J, Syn N, Foo R, Khoo CM, Wang JW, Dimitriadis GK, Young DY, Siddiqui MS, Lam CSP, Wang Y, Figtree GA, Chan MY, Cummings DE, Noureddin M, Wong VWS, Ma RCW, Mantzoros CS, Sanyal A, Muthiah MD. The global burden of metabolic disease: Data from 2000 to 2019. Cell Metab. 2023;35(3):414–428.e3.

2. Noubiap JJ, Nansseu JR, Lontchi-Yimagou E, Nkeck JR, Nyaga UF, Ngouo AT, Tounouga DN, Tianyi FL, Foka AJ, Ndoadoumgue AL, Bigna JJ. Geographic distribution of metabolic syndrome and its components in the general adult population: A meta-analysis of global data from 28 million individuals. Diabetes Res. Clin. Pract. 2022;188. doi:10.1016/j.diabres.2022.109924.

3. Saklayen MG. The Global Epidemic of the Metabolic Syndrome. Curr. Hypertens. Rep. 2018;20(2). doi:10.1007/S11906-018-0812-Z.

4. Silveira Rossi JL, Barbalho SM, Reverete de Araujo R, Bechara MD, Sloan KP, Sloan LA. Metabolic syndrome and cardiovascular diseases: Going beyond traditional risk factors. Diabetes. Metab. Res. Rev. 2022;38(3). doi:10.1002/DMRR.3502.

5. McHill AW, Wright KP. Role of sleep and circadian disruption on energy expenditure and in metabolic predisposition to human obesity and metabolic disease. Obesity Reviews 2017;18:15–24.

6. Hostinar CE, Ross KM, Chen E, Miller GE. Early-life socioeconomic disadvantage and metabolic health disparities. Psychosom. Med. 2017;79(5):514–523.

7. Stefanaki C, Pervanidou P, Boschiero D, Chrousos GP. Chronic stress and body composition disorders: implications for health and disease. Hormones 2018;17(1):33–43.

8. Picard M, Juster RP, McEwen BS. Mitochondrial allostatic load puts the “gluc” back in glucocorticoids. Nat. Rev. Endocrinol. 2014;10(5):303–310.

9. Herman JP, Mcklveen JM, Ghosal S, Kopp B, Wulsin A, Makinson R, Scheimann J, Myers B. Regulation of the hypothalamic-pituitary-adrenocortical stress response. Compr. Physiol. 2016;6(2).

10. Chrousos GP. The concepts of stress and stress system disorders. Overview of physical and behavioral homeostasis. JAMA: The Journal of the American Medical Association 1992;267(9):1244–1252.

11. Myers B, McKlveen JM, Herman JP. Glucocorticoid actions on synapses, circuits, and behavior: Implications for the energetics of stress. Front. Neuroendocrinol. 2014;35(2). doi:10.1016/j.yfrne.2013.12.003.

12. Dearing C, Handa RJ, Myers B. Sex differences in autonomic responses to stress: implications for cardiometabolic physiology. Am. J. Physiol. Endocrinol. Metab. 2022;323(3). doi:10.1152/ajpendo.00058.2022.

13. Contreras-Zentella ML, Hernández-Muñoz R. Possible gender influence in the mechanisms underlying the oxidative stress, inflammatory response, and the metabolic alterations in patients with obesity and/or type 2 diabetes. Antioxidants 2021;10(11). doi:10.3390/ANTIOX10111729,

14. McEwen BS. Central effects of stress hormones in health and disease: Understanding the protective and damaging effects of stress and stress mediators. Eur. J. Pharmacol. 2008;583(2–3):174–185.

15. Pervanidou P, Chrousos GP. Stress and obesity/metabolic syndrome in childhood and adolescence. International Journal of Pediatric Obesity 2011;6(SUPPL. 1):21–28.

16. Merz EC, Myers B, Hansen M, Simon KR, Strack J, Noble KG. Socioeconomic Disparities in Hypothalamic-Pituitary-Adrenal Axis Regulation and Prefrontal Cortical Structure. Biological Psychiatry Global Open Science 2024;4(1). doi:10.1016/j.bpsgos.2023.10.004.

17. Dearing C, Morano R, Ptaskiewicz E, Mahbod P, Scheimann JR, Franco-Villanueva A, Wulsin L, Myers B. Glucoregulation and coping behavior after chronic stress in rats: Sex differences across the lifespan. Horm. Behav. 2021;136. doi:10.1016/j.yhbeh.2021.105060.

18. Hamer M, Steptoe A. Cortisol responses to mental stress and incident hypertension in healthy men and women. Journal of Clinical Endocrinology and Metabolism 2012;97(1). doi:10.1210/JC.2011-2132,

19. Yusuf PS, Hawken S, Ôunpuu S, Dans T, Avezum A, Lanas F, McQueen M, Budaj A, Pais P, Varigos J, Lisheng L. Effect of potentially modifiable risk factors associated with myocardial infarction in 52 countries (the INTERHEART study): case-control study. Lancet 2004;364(9438):937–952.

20. Blackburn-Munro G, Blackburn-Munro RE. Chronic pain, chronic stress and depression: Coincidence or consequence? J. Neuroendocrinol. 2001;13(12):1009–1023.

21. Pace TWW, Mletzko TC, Alagbe O, Musselman DL, Nemeroff CB, Miller AH, Heim CM. Increased stress-induced inflammatory responses in male patients with major depression and increased early life stress. American Journal of Psychiatry 2006;163(9):1630–1633.

22. Cohen S, Schwartz JE, Epel E, Kirschbaum C, Sidney S, Seeman T. Socioeconomic status, race, and diurnal cortisol decline in the Coronary Artery Risk Development in Young Adults (CARDIA) Study. Psychosom. Med. 2006;68(1):41–50.

23. Myers B, McKlveen JM, Morano R, Ulrich-Lai YM, Solomon MB, Wilson SP, Herman JP. Vesicular glutamate transporter 1 knockdown in infralimbic prefrontal cortex augments neuroendocrine responses to chronic stress in male rats. Endocrinology 2017;158(10). doi:10.1210/en.2017-00426.

24. Pace SA, Christensen C, Schackmuth MK, Wallace T, McKlveen JM, Beischel W, Morano R, Scheimann JR, Wilson SP, Herman JP, Myers B. Infralimbic cortical glutamate output is necessary for the neural and behavioral consequences of chronic stress. Neurobiol. Stress 2020;13. doi:10.1016/j.ynstr.2020.100274.

25. Wallace T, Schaeuble D, Pace SA, Schackmuth MK, Hentges ST, Chicco AJ, Myers B. Sexually divergent cortical control of affective-autonomic integration. Psychoneuroendocrinology 2021;129. doi:10.1016/J.PSYNEUEN.2021.105238.

26. Wood M, Adil O, Wallace T, Fourman S, Wilson SP, Herman JP, Myers B. Infralimbic prefrontal cortex structural and functional connectivity with the limbic forebrain: a combined viral genetic and optogenetic analysis. Brain Struct. Funct. 2019;224(1). doi:10.1007/s00429-018-1762-6.

27. Lukinic E, Wallace T, McCartney C, Myers B. Infralimbic prefrontal cortical projections to the autonomic brainstem: quantification of inputs to cholinergic and adrenergic/noradrenergic nuclei. Brain Struct. Funct. 2025;230(7):117.

28. Kim A, Knudsen JG, Madara JC, Benrick A, Hill T, Kadir LA, Kellard JA, Mellander L, Miranda C, Lin H, James T, Suba K, Spigelman AF, Wu Y, Macdonald PE, Asterholm IW, Magnussen T, Christensen M, Visboll T, Salem V, Knop FK, Rorsman P, Lowell BB, Briant LJB. Arginine-vasopressin mediates counter-regulatory glucagon release and is diminished in type 1 diabetes. Elife 2021;10. doi:10.7554/ELIFE.72919,

29. Pace SA, Myers B. Hindbrain Adrenergic/Noradrenergic Control of Integrated Endocrine and Autonomic Stress Responses. Endocrinology (United States) 2024;165(1). doi:10.1210/endocr/bqad178.

30. Li AJ, Wang Q, Ritter S. Selective pharmacogenetic activation of catecholamine subgroups in the ventrolateral medulla elicits key glucoregulatory responses. Endocrinology 2018;159(1):341–355.

31. Guyenet PG, Stornetta RL, Bochorishvili G, DePuy SD, Burke PGR, Abbott SBG. C1 neurons: the body’s EMTs. Am. J. Physiol. Regul. Integr. Comp. Physiol. 2013;305(3). doi:10.1152/AJPREGU.00054.2013.

32. Zhao Z, Wang L, Gao W, Hu F, Zhang J, Ren Y, Lin R, Feng Q, Cheng M, Ju D, Chi Q, Wang D, Song S, Luo M, Zhan C. A Central Catecholaminergic Circuit Controls Blood Glucose Levels during Stress. Neuron 2017;95(1):138–152.e5.

33. Zsombok A, Desmoulins LD, Derbenev A V. Sympathetic circuits regulating hepatic glucose metabolism: where we stand. Physiol. Rev. 2024;104(1):85–101.

34. Pace SA, Lukinic E, Wallace T, McCartney C, Myers B. Cortical–brainstem circuitry attenuates physiological stress reactivity. Journal of Physiology 2024;602(5). doi:10.1113/JP285627.

35. Sando R, Bushong E, Zhu Y, Huang M, Considine C, Phan S, Ju S, Uytiepo M, Ellisman M, Maximov A. Assembly of Excitatory Synapses in the Absence of Glutamatergic Neurotransmission. Neuron 2017;94(2):312–321.e3.

36. Schaeuble D, Wallace T, Pace SA, Hentges ST, Myers B. Sex-specific prefrontal-hypothalamic control of behavior and stress responding. Psychoneuroendocrinology 2024;159. doi:10.1016/j.psyneuen.2023.106413.

37. Link E, Edelmann L, Chou JH, Binz T, Yamasaki S, Eisel U, Baumert M, Südhof TC, Niemann H, Jahn R. Tetanus toxin action: inhibition of neurotransmitter release linked to synaptobrevin proteolysis. Biochem. Biophys. Res. Commun. 1992;189(2):1017–1023.

38. Myers B, McKlveen JM, Morano R, Ulrich-Lai YM, Solomon MB, Wilson SP, Herman JP. Vesicular glutamate transporter 1 knockdown in infralimbic prefrontal cortex augments neuroendocrine responses to chronic stress in male rats. Endocrinology 2017;158(10):3579–3591.

39. Schaeuble D, Packard AEB, McKlveen JM, Morano R, Fourman S, Smith BL, Scheimann JR, Packard BA, Wilson SP, James J, Hui DY, Ulrich-Lai YM, Herman JP, Myers B. Prefrontal Cortex Regulates Chronic Stress-Induced Cardiovascular Susceptibility. J. Am. Heart Assoc. 2019;8(24):e014451.

40. Dearing C, Sanford E, Olmstead N, Morano R, Wulsin L, Myers B. Sex-specific cardiac remodeling in aged rats after adolescent chronic stress: associations with endocrine and metabolic factors. Biol. Sex Differ. 2024;15(1). doi:10.1186/S13293-024-00639-7.

41. Ghosal S, Nunley A, Mahbod P, Lewis AG, Smith EP, Tong J, D’Alessio DA, Herman JP. Mouse handling limits the impact of stress on metabolic endpoints. Physiol. Behav. 2015;150:31–37.

42. Winarto H, Habiburrahman M, Febriana IS, Kusuma F, Nuryanto KH, Anggraeni TD, Utami TW, Putra AD. Is there any difference in insulin resistance status between cases of benign and malignant ovarian neoplasms? A study on surrogate markers of insulin resistance in Indonesian non-diabetic women. Oncol. Lett. 2022;25(1). doi:10.3892/OL.2022.13609.

43. Mather K. Surrogate measures of insulin resistance: of rats, mice, and men. Am. J. Physiol. Endocrinol. Metab. 2009;296(2). doi:10.1152/AJPENDO.90889.2008.

44. González-González JG, Violante-Cumpa JR, Zambrano-Lucio M, Burciaga-Jimenez E, Castillo-Morales PL, Garcia-Campa M, Solis RC, González-Colmenero AD, Rodríguez-Gutiérrez R. HOMA-IR as a predictor of Health Outcomes in Patients with Metabolic Risk Factors: A Systematic Review and Meta-analysis. High Blood Press. Cardiovasc. Prev. 2022;29(6):547–564.

45. Packard AEB, Ghosal S, Herman JP, Woods SC, Ulrich-Lai YM. Chronic variable stress improves glucose tolerance in rats with sucrose-induced prediabetes. Psychoneuroendocrinology 2014;47:178–188.

46. Moh Moh MA, Jung CH, Lee B, Choi D, Kim BY, Kim CH, Kang SK, Mok JO. Association of glucagon-to-insulin ratio and nonalcoholic fatty liver disease in patients with type 2 diabetes mellitus. Diab. Vasc. Dis. Res. 2019;16(2):186–195.

47. Unger RH. Glucagon and the insulin: glucagon ratio in diabetes and other catabolic illnesses. Diabetes 1971;20(12):834–838.

48. Kalra S, Gupta Y. The Insulin:Glucagon Ratio and the Choice of Glucose-Lowering Drugs. Diabetes Ther. 2016;7(1):1–9.

49. Desmoulins LD, Molinas AJR, Dugas CM, Williams GL, Kamenetsky S, Davis RK, Derbenev A V., Zsombok A. A subset of neurons in the paraventricular nucleus of the hypothalamus directly project to liver-related premotor neurons in the ventrolateral medulla. Auton. Neurosci. 2025;257. doi:10.1016/j.autneu.2024.103222.

50. Mauvais-Jarvis F. Sex differences in metabolic homeostasis, diabetes, and obesity. Biology of Sex Differences 2015 6:1 2015;6(1):14-.

51. Gabbott PLA, Warner T, Busby SJ. Catecholaminergic neurons in medullary nuclei are among the post-synaptic targets of descending projections from infralimbic area 25 of the rat medial prefrontal cortex. Neuroscience 2007;144(2):623–635.

52. Goldstein JM, Langer A, Lesser JA. Sex Differences in Disorders of the Brain and Heart-A Global Crisis of Multimorbidity and Novel Opportunity. JAMA Psychiatry 2021;78(1):7–8.

53. Goldstein JM, Hale T, Foster SL, Tobet SA, Handa RJ. Sex differences in major depression and comorbidity of cardiometabolic disorders: impact of prenatal stress and immune exposures. Neuropsychopharmacology 2019;44(1):59–70.

54. Bale TL, Epperson CN. Sex differences and stress across the lifespan. Nat. Neurosci. 2015;18(10):1413–1420.

55. Mauvais-Jarvis F. Sex differences in energy metabolism: natural selection, mechanisms and consequences. Nat. Rev. Nephrol. 2024;20(1):56–69.

56. Mauvais-Jarvis F, Lindsey SH. Metabolic benefits afforded by estradiol and testosterone in both sexes: clinical considerations. J. Clin. Invest. 2024;134(17). doi:10.1172/JCI180073.

57. Yan H, Yang W, Zhou F, Li X, Pan Q, Shen Z, Han G, Newell-Fugate A, Tian Y, Majeti R, Liu W, Xu Y, Wu C, Allred K, Allred C, Sun Y, Guo S. Estrogen Improves Insulin Sensitivity and Suppresses Gluconeogenesis via the Transcription Factor Foxo1. Diabetes 2019;68(2):291–304.

58. Wada T, Hori S, Sugiyama M, Fujisawa E, Nakano T, Tsuneki H, Nagira K, Saito S, Sasaoka T. Progesterone inhibits glucose uptake by affecting diverse steps of insulin signaling in 3T3-L1 adipocytes. 2010;298(4). doi:10.1152/ajpendo.00649.2009.

59. Pitteloud N, Mootha VK, Dwyer AA, Hardin M, Lee H, Eriksson KF, Tripathy D, Yialamas M, Groop L, Elahi D, Hayes FJ. Relationship between testosterone levels, insulin sensitivity, and mitochondrial function in men. Diabetes Care 2005;28(7):1636–1642.

60. Brivio E, Kos A, Ulivi AF, Karamihalev S, Ressle A, Stoffel R, Hirsch D, Stelzer G, Schmidt M V., Lopez JP, Chen A. Sex shapes cell-type-specific transcriptional signatures of stress exposure in the mouse hypothalamus. Cell Rep. 2023;42(8):112874.

61. Brivio E, Lopez JP, Chen A. Sex differences: Transcriptional signatures of stress exposure in male and female brains. Genes Brain Behav. 2020;19(3):e12643.

62. Moore SR, Halldorsdottir T, Martins J, Lucae S, Müller-Myhsok B, Müller NS, Piechaczek C, Feldmann L, Freisleder FJ, Greimel E, Schulte-Körne G, Binder EB, Arloth J. Sex differences in the genetic regulation of the blood transcriptome response to glucocorticoid receptor activation. Transl. Psychiatry 2021;11(1). doi:10.1038/s41398-021-01756-2.

63. Zhang Y, Klein K, Sugathan A, Nassery N, Dombkowski A, Zanger UM, Waxman DJ. Transcriptional Profiling of Human Liver Identifies Sex-Biased Genes Associated with Polygenic Dyslipidemia and Coronary Artery Disease. PLoS One 2011;6(8):e23506.

64. Van Nas A, Guhathakurta D, Wang SS, Yehya N, Horvath S, Zhang B, Ingram-Drake L, Chaudhuri G, Schadt EE, Drake TA, Arnold AP, Lusis AJ. Elucidating the role of gonadal hormones in sexually dimorphic gene coexpression networks. Endocrinology 2009;150(3):1235–1249.

